# Target of rapamycin (TOR) regulates *CURLY LEAF* (*CLF*) translation in response to environmental stimuli

**DOI:** 10.1101/2024.06.28.601127

**Authors:** Yihan Dong, Ez-Zahra Oubassou, Elise Hoffmann, Wenna Zhang, Veli Uslu, Alexandre Berr

**Affiliations:** Institute de Biologie Moléculaire des Plantes, Centre National de la Recherche Scientifique, UPR 2357, Université de Strasbourg, Strasbourg, France; Cadi Ayyad University, Faculty of Sciences Semlalia, Laboratory of Microbial Biotechnologies, Agrosciences and Environment (BioMAgE), Marrakech, Morocco; Beijing Key Laboratory of Growth and Developmental Regulation for Protected Vegetable Crops, China Agricultural University, Beijing, 100193, China; RLP AgroScience GmbH, Neustadt an der Weinstraße, 67435, Germany; MAPS, Center for Organismal Studies, Heidelberg University, Heidelberg, 69120, Germany

**Keywords:** Target of rapamycin, translation reinitiation, stress response, chromatin, polycomb repressive complex

## Abstract

The target of rapamycin (TOR)-Polycomb repressive complex 2 (PRC2) pathway is a crucial link that translates environmental and developmental cues into chromatin, thus reprogramming transcription. While the PRC2 methyltransferase Curly leaf (CLF) is known to be specifically involved, the underlying mechanism remains unclear. This study sheds light on how TOR fine-tunes CLF protein levels by promoting translation re-initiation mediated by eIF3h. We found that the second upstream open reading frame (uORF) located in the 5’ leader region of the *CLF* transcript significantly represses its translation (by 50%). Plants lacking this uORF leader displayed reduced sensitivity to TOR inhibition and impaired induction of stress-responsive genes. Interestingly, this uORF sequence exhibits partial conservation across diverse plant species, suggesting a potential role in adaptation to various environmental conditions. Our findings reveal a dynamic mechanism within the TOR-PRC2 pathway, highlighting its responsiveness to environmental stimuli.

## Introduction

Plants, as sessile organisms, continuously adjust their developmental programs to cope with the ever-changing environment. This adaptation is intricately regulated by signaling pathways, including those involving the Target of Rapamycin (TOR) and Polycomb Repressive Complex 2 (PRC2). TOR, acting as a sensor kinase, governs protein synthesis, while PRC2 catalyzes H3 lysine 27 tri-methylation (H3K27me3) to modulate gene expression. Both of these pathways play crucial roles in integrating environmental signals to orchestrate developmental responses. In mammals, TOR operates within two functionally and structurally distinct protein complexes ^1^. However, it is noteworthy that only TOR complex 1 is conserved in plants, a homodimer complex composed of mLST8 (mammalian Lethal with Sec13 protein 8) and Raptor (Regulatory-Associated Protein of mTOR) ^2^. PRC2, on the other hand, is evolutionarily conserved and comprises four core protein subunits across eukaryotes. Arabidopsis possesses three orthologs of the H3K27 trimethyltransferase Enhancer of zeste (E(z)), namely CURLY LEAF (CLF), SWINGER (SWN), and MEDEA (MEA), three orthologs of Su(z)12, named EMBRYONIC FLOWER 2 (EMF2), VERNALIZATION 2 (VRN2), and FERTILIZATION-INDEPENDENT SEED 2 (FIS2), which stabilize PRC2, one ortholog of Esc named FERTILIZATION-INDEPENDENT ENDOSPERM (FIE), and five orthologs of the nucleosome remodeling factor Nurf55 termed MULTICOPY SUPPRESSOR OF IRA 1–5 (MSI1-5) ^3^.

The TOR-PRC2 pathway has gained significant attention in animals. Specifically, mTORC1 activates its canonical substrate S6K1, leading to phosphorylation of histone H2B and the promotion of H3K27me3 at genes linked to adipogenesis ^4^. Recent findings have expanded our understanding, revealing that both mTORC1 and mTORC2 play roles in regulating H3K27me3. In human glioblastoma tumor cells, mTORC1 influences the translation of the E(Z) homolog EZH2, while mTORC2 affects the metabolism of S-adenosylmethionine (SAM), the principal methyl donor for histone methylation reactions ^5^. In mice, a mutation in Raptor, a component of mTORC1, reduces EED levels (the mammalian ortholog of FIE), suggesting a link between mTORC1 activity and PRC2 production ^6^. Additionally, inhibition of TOR by rapamycin increases the translation of H3/H4 in Drosophila and mice, causing chromatin rearrangement and heterochromatin relocation ^7^. Together, these findings hint towards the intricate role of TOR-mediated translation control in the regulation of H3K27me3 deposition in animals.

Translation, the most energy-intensive cellular process, undergoes intricate regulation. In mammals, mTOR orchestrates translation initiation by phosphorylating two primary substrates: eIF4E-binding proteins (4EBPs) and ribosomal protein S6 kinases (S6Ks) ^8^. In plants, TOR plays a prominent role in the translation reinitiation of numerous mRNAs harboring upstream open reading frames (uORFs) within their 5’ leader regions. Typically, following translation termination, the 80S complex disassembles and faces challenges in reinitiating translation at a downstream ORF on the same mRNA. Both mammals and plants allow reinitiation only if the first uORF is short. Conversely, a relatively long uORF (>16 codons) or a combination of several short uORFs can impede the resumption of scanning and the initiation of the downstream main ORF by post-terminating ribosomes, thereby reducing mRNA translation efficiency by 30-80% ^9, 10^. In plants, an active TOR relay can overcome uORF repression by promoting reinitiation after uORF translation through a series of phosphorylation events. This process includes the phosphorylation of eIF3 subunit h (eIF3h), reinitiation-supporting protein (RISP), and eukaryotic ribosomal protein S6 (eS6) ^11, 12^.

In plants, the connection between TOR and PRC2 has been elucidated from two distinct but related perspectives. In a developmental context, TOR directly phosphorylates the unique PRC2 component FIE, facilitating its translocation to the nucleus and H3K27me3 deposition at crucial developmental genes ^13^. In the more dynamic process of stress response, we detailed a TOR-PRC2 pathway specifically involving the histone H3K27 tri-methyltransferase CLF ^14^. Genes repressed by TOR were found to be preferentially associated with either bistable or silent chromatin states. Interestingly, while both states exhibit a high level of H3K27me3, the bistable state is additionally marked by an elevated level of the active histone mark H3K4me3 ^15^. Interestingly, TOR was revealed to regulate H3K4me3, likely through the SWI/SNF-related chromatin remodeler BRAHMA (BRM) ^16, 17^ and the H3K4 tri-methyltransferase ARABIDOPSIS TRITHORAX 1 (ATX1) ^18^. Together, these findings suggest two distinct groups of TOR-repressed genes with varying chromatin accessibility, responsiveness, and functionality. Furthermore, while TOR inhibition results in a substantial global reduction of H3K27me3, other marks such as H3K4me3, H3K9me2, and H3K36me3 appear unaffected ^13^.

While H3K27me3 is a pivotal histone mark regulated by the TOR pathway, the specific requirement of CLF within the TOR-PRC2 pathway, as previously reported ^14^, remains elusive. In this study, we delve into the regulatory mechanisms connecting TOR with CLF, unveiling that TOR governs H3K27me3 loading via translation reinitiation of the uORF-containing *CLF* mRNA. The abundance of CLF protein and its responsiveness to TOR activity emerged as crucial determinants in plant stress response, shedding light on its potential role in environmental adaptability. Our findings provide new insights into how TOR might orchestrate stress by modulating CLF-PRC2 assembly dynamics.

## Results

### TOR promotes translation reinitiation to overcome the repressive uORF in the *CLF* leader

While the role of PRC2 in plant development and the evolution of its different subunit have been extensively examined ^3^, little attention has been given to how these PRC2 components are produced and assembled. Upon meticulous examination of mRNAs encoding *CLF* and *SWN*, a distinctive feature emerged: only the *CLF* mRNA contains two upstream open reading frames (uORFs) within its 5’UTR, which have a high predicted inhibition potential (**Fig. 1A**). This prediction was based on the uORF length (> 16 codons) and an optimal initiation context of AUG. Notably, uORFs with AUGs residing in a weak initiation context, lacking both R (A or G) at position -3 and G at position +4 relative to the first nucleotide of the start codon, are typically associated with skipped recognitions ^10^.

**Figure 1:**
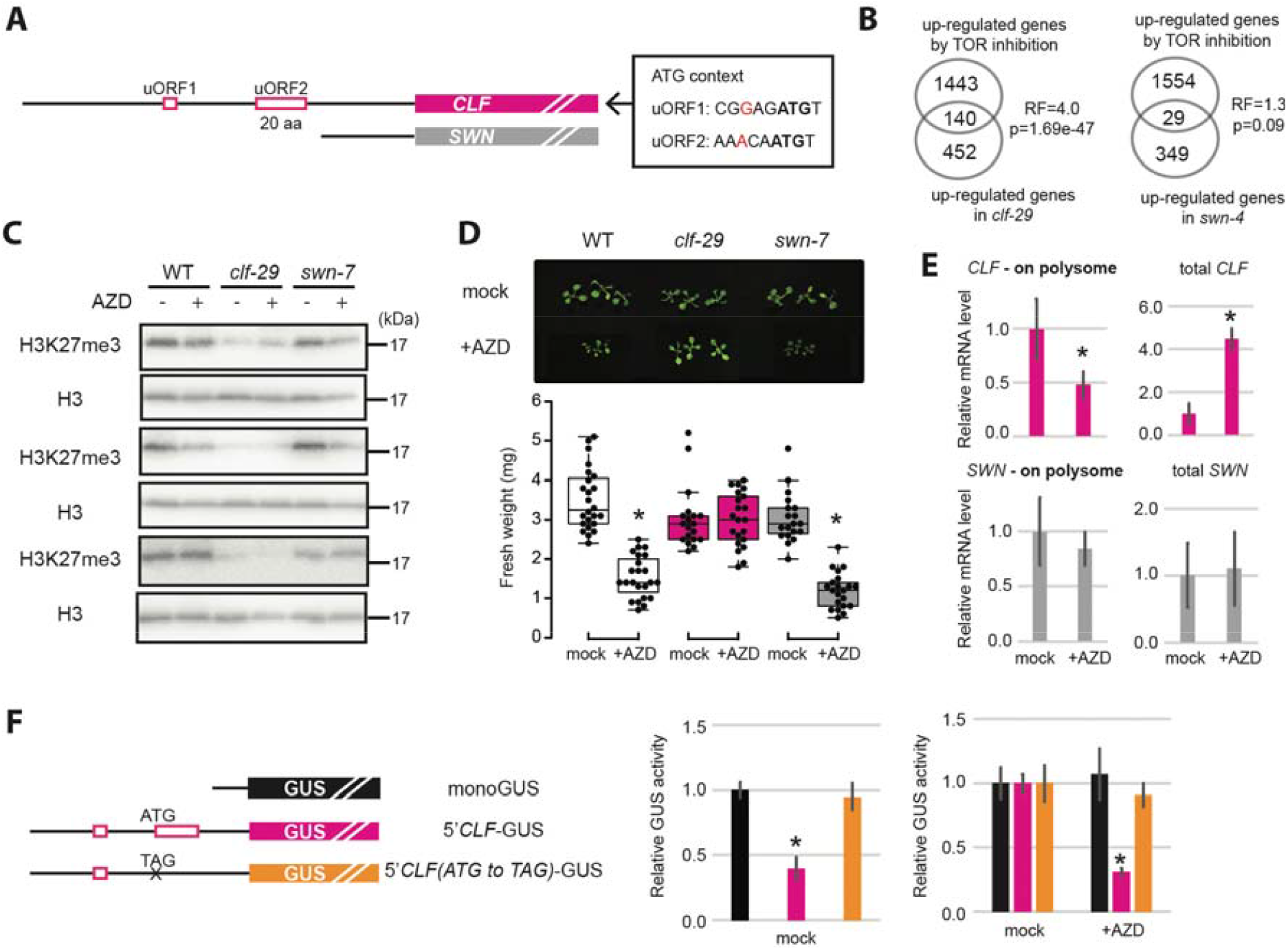
TOR regulates CLF translation via the 2nd uORF in its 5’UTR. (A) Schematic representation of 5 ‘UTR features from *CLF* and *SWN* mRNAs. Red boxes represent putative inhibitory uORFs with optimal AUG initiation context. All 5 ‘UTR sequences were obtained from JBrowse (TAIR10 genome). (B) Overlap of TOR-repressed genes with CLF/SWN-repressed genes (hypergeometric test, RF: representation factor, RF > 1 indicates more overlap than expected, \**p* < 0.05). TOR-repressed genes and CLF/SWN-repressed genes were extracted from Dong et al., (2015) and Shu et al., (2019) (fold change > 2 and *p* < 0.05). (C) Western blot quantification of H3K27 me3 levels in *clf-29* and *swn-7* upon TOR inhibition by AZD8055. (D) Fresh weight of Arabidopsis seedlings grown for 1 week on medium supplemented with or without 0.5 μ M AZD8055. Shoot fresh weight was measured under mock and AZD8055-applied conditions and ratio was calculated to indicate the growth sensitivity to TOR inhibition (n > 20, mean ± s.d., \**p* < 0.05). (E) Quantification of total *CLF* and *SWN* mRNA and polysome loading upon TOR inhibition by 1 μ M AZD8055 for 16 hours (n = 3, mean ± s.d., \**p* < 0.05). mRNA level associated with polysomes reflects the translation efficiency. (F) Mutation of the AUG start codon in the 2nd uORF of CLF leader fused to a GUS reporter. The reporter construct was transfected into protoplast and GUS activity was measured to evaluate uORF impact on main ORF translation. Values are presented relative to the monoGUS construct (left histogram) or relative to the mock treatment for each individual construct (right histogram). 0 .2 μM AZD8055 was applied for 16 hours (n = 3, mean ± s.d., different letters indicate *p* < 0.05).

To further explore the distinctive role of CLF, as opposed to SWN, in the TOR-PRC2 pathway, we compared first the transcriptomic responses to TOR inhibition ^19^ or *CLF*/*SWN* knockout ^20^. Strikingly, TOR-responsive genes exhibited a pronounced overlap with up-regulated genes in the *clf-29* mutant, but not in the *swn* mutant (**Fig. 1B**). Additionally, treatment with the TOR inhibitor AZD8055 significantly reduced H3K27me3 levels in both wild-type and *swn* mutant, contrasting with the *clf-29* mutant where this decrease was not observed (**Fig. 1C**). Furthermore, growth sensitivity to TOR inhibition revealed that while the *swn* mutant demonstrated similar AZD sensitivity to the wild type, *clf-29*, as previously reported ^14^, exhibited marked insensitivity to TOR inhibition (**Fig. 1D**).

Recent high-resolution ribosome profiling, notably depicted in the translated uORFs (TuORFs) track on JBrowse ^21^, has unveiled a myriad of previously unknown translation events on uORFs, downstream ORFs, and small RNAs. Notably, the second uORF in the 5’UTR of *CLF* (**Fig. 1A**) exhibited ribosome occupancy, significantly enhancing its likelihood of a repressive effect and highlighting the potential role of TOR in regulating *CLF* translation. To delve deeper into this, we investigated the translation efficiency of *CLF* using ribosome profiling. Intriguingly, upon TOR inhibition, less *CLF* mRNA was associated with actively translating polyribosomes, whereas *SWN* mRNAs remained unaffected (**Fig. 1E**). These findings collectively suggest that CLF translation is TOR-dependent, while SWN translation is TOR-insensitive. To further substantiate the regulatory role of TOR on CLF translation, we conducted experiments using Arabidopsis protoplasts. By expressing the *CLF* leader together with either a wild-type or a mutated version of the second uORF fused to a GUS reporter. Resulting GUS activities demonstrated that the wild-type second uORF functions as a repressor, and this repressive effect is augmented upon TOR inhibition (**Fig. 1F**). Altogether, these results underscore that CLF is primarily responsible for the TOR signaling output, likely owing to translational control by TOR.

TOR-eIF3h axis is the most established to regulate plant translation reinitiation after uORFs ^12^. Therefore, we analyzed the *CLF* and *SWN* mRNA loading on polysomes in *eif3h1* mutant. Upon eIF3h knockout, *CLF* mRNA showed a notable shift from heavy polysome to light polysome and 80S ribosome, while *SWN* remained relatively unaffected (**Fig. 2A**). This polysome loading pattern suggest that CLF translation is finetuned by its uORF rather than being completely inhibited. Under normal growth conditions, TOR activity and reinitiation capacity are maintained at a housekeeping level and eIF3h mutation did not visibly alter global H3K27me3 level. However, upon TOR hyperactivation by CaMV infection ^22^, *eif3h1* mutant failed to establish proper H3K27me3 (**Fig. 2B**).

**Figure 2:**
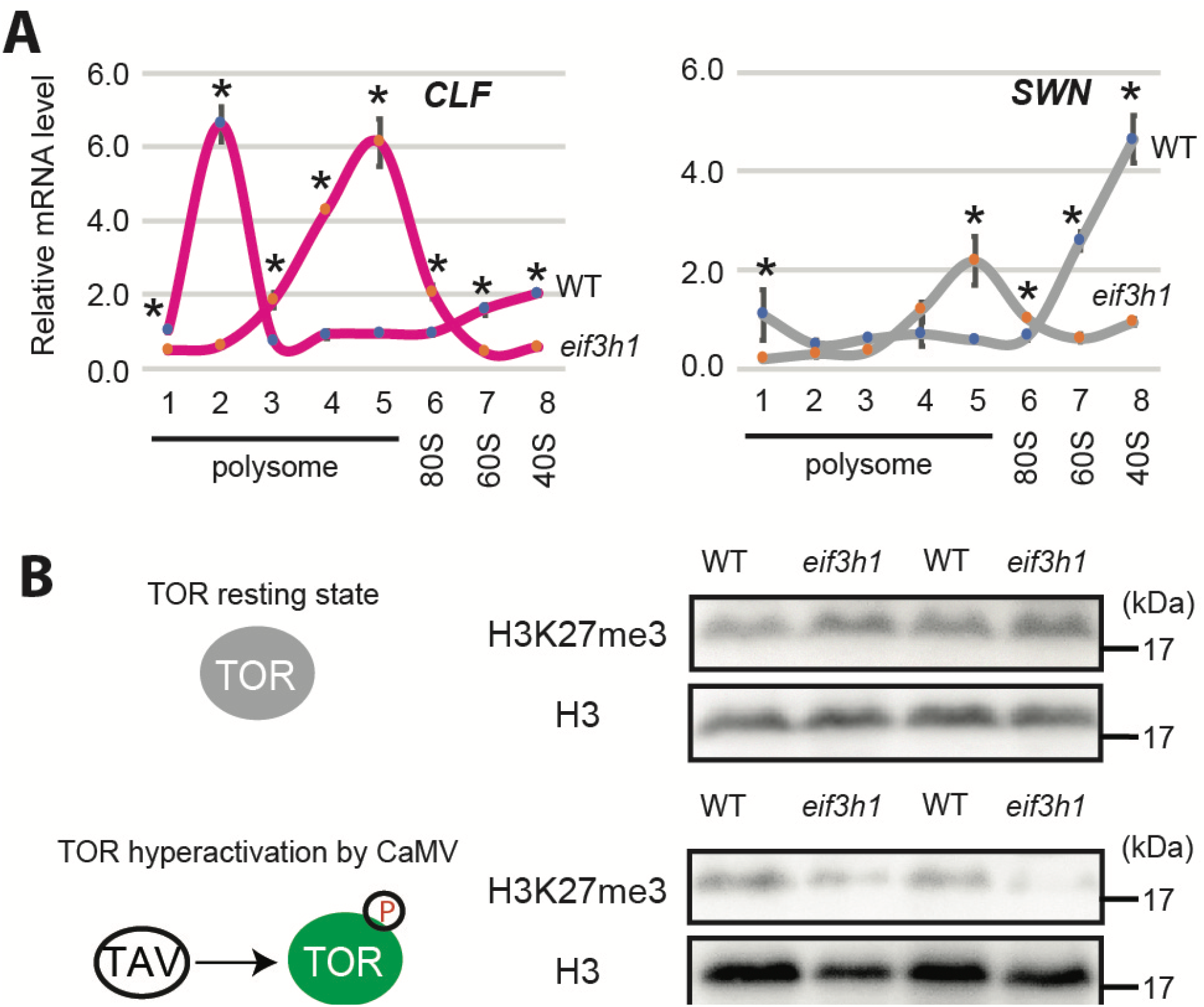
TOR regulates CLF translation via eIF3 h. (A) Quantification of *CLF* and *SWN* mRNA in different ribosome fractions, including polysomes, monosome (80S), and large (60S) and small (40S) ribosome subunits, in the *eif3h1* mutant (n = 3, mean ± s.d., \**p* < 0 .05). (B) H3 K27 me3 levels in the *eif3h1* mutant under normal growth conditions and TOR hyperactivation. TOR hyperactivation was induced by Cauliflower mosaic virus (CaMV) infection. Samples were collected at 20 days post-inoculation. TAV, CaMV trans-activator protein.

### 5’UTR of *CLF* mediates its translational control by TOR

To unravel functional relevance of *CLF*’s uORF-containing 5’UTR, we used various CLF-GFP reporter lines. Our primary focus was on the *clf-29* mutant complemented with the CLF genomic sequence (*clf-29* CLF-gDNA) ^23^. For comparison, we also used a *swn* mutant in the *clf-29* CLF-gDNA background (*clf swn* CLF-gDNA) and compared it with the wild type Col-0. Additionally, a CLF-GFP line without 5’UTR (**Supplementary figure 1**), in the *clf-50* mutant background (*clf-50* CLF-CDS) ^24^, was compared to its wild type WS.

Subsequently, we analyzed the sensitivity to TOR inhibition in the different CLF complementation lines with or without 5’UTRs. In comparison to both *clf-29* CLF-gDNA and *clf swn* CLF-gDNA lines, the *clf-50* CLF-CDS line displayed a delayed response to TOR inhibition (**Fig. 3A**). Using the GFP reporter, we conducted a comprehensive analysis of CLF protein levels through confocal microscopy and western blotting. In the *clf-29* CLF-gDNA line, the protein level of CLF rapidly declined upon TOR inhibition (**Fig. 3B, C**). Conversely, the CLF protein level in the *clf-50* CLF-CDS line exhibited resistance to TOR inhibition (**Fig. 3B, C**). The elevated CLF protein level in the *clf swn* CLF-gDNA line (**Fig. 3C**) is likely due to the increased *CLF* transcript level (**Supplementary figure 2**), while it maintained responsiveness to TOR inhibition. Taken together, these findings strongly suggest that TOR exerts control over CLF translation, with the 5’UTR playing a dominant role in mediating this regulatory response.

**Figure 3:**
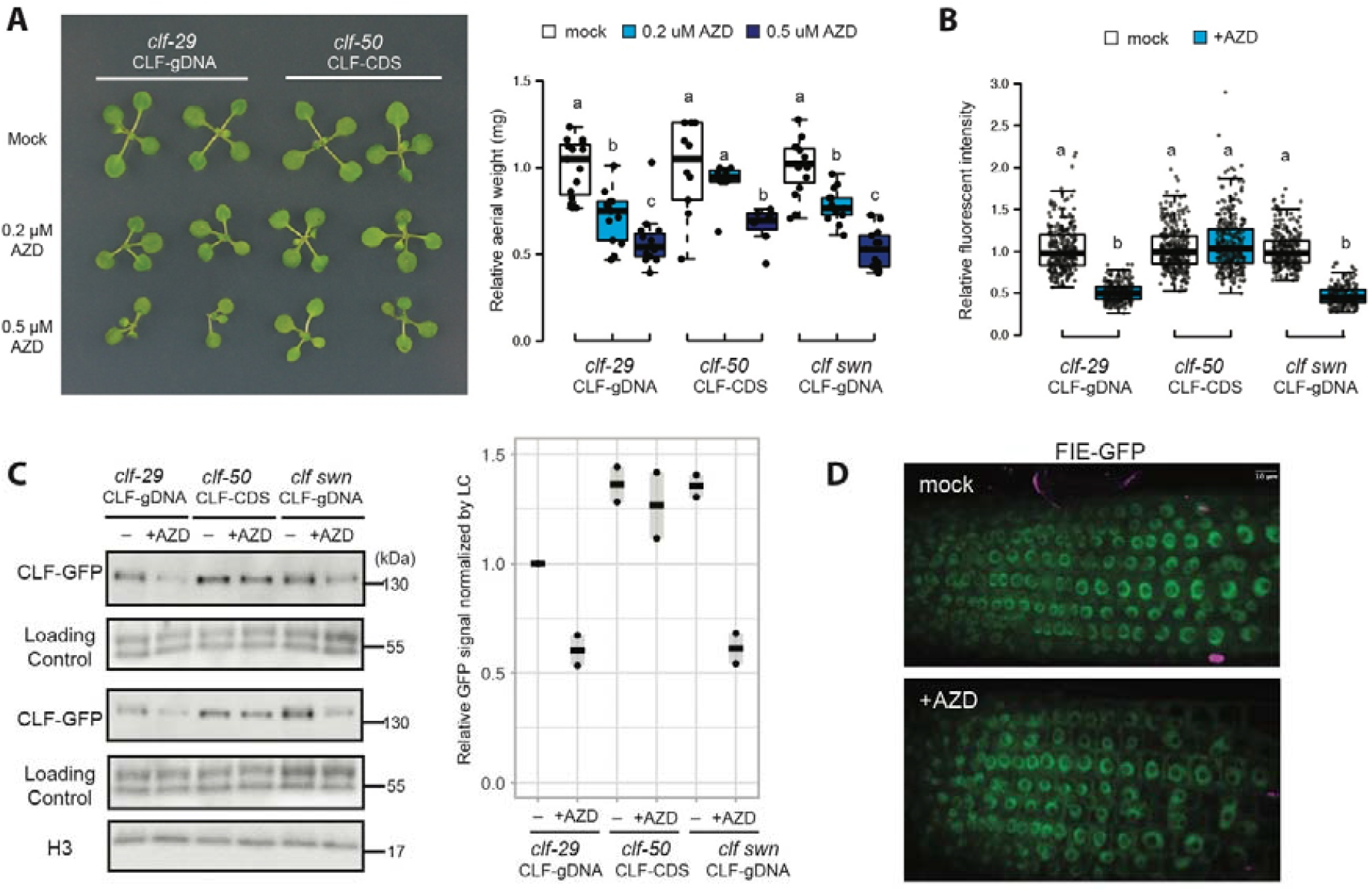
CLF 5’UTR mediates translational control by TOR. (A) Arabidopsis seedlings were germinated on 1/2MS medium for 4 days and then transferred to medium supplemented with or without 0.2/0.5 μM AZD8055 for an additional 7 days (One-way ANOVA, different letters indicate significant differences, *p* < 0.05). (B, C) CLF protein levels were quantified by confocal microscope (B) and western blot (C; results from two biological replicates are presented) in *clf* mutants complemented with CLF genomic sequence (CLF-gDNA) or coding domain sequence (CLF-CDS) following TOR inhibition with 1 μ M AZD8055 for 16 h. The *swn-4* (in the background of *clf-29* CLF-gDNA) mutant was used as a control (LC = Loading Control). (D) FIE-GFP line was treated with 1 μM TOR kinase inhibitor AZD8055 for 16 h.

Combining our findings with those from Ye et al. (2022), it appears that TOR regulates PRC2 through mechanisms involving CLF translation and FIE phosphorylation. The biological significance of the TOR-FIE pathway, primarily explored in the context of vernalization, involved the cytoplasmic retention of FIE in response to strong TOR inhibition (*i*.*e*., 10 μM AZD8055 for 3 days) ^13^. The TOR-FIE pathway knockout line exhibits severe developmental defects, underscoring the critical role of FIE phosphorylation by TOR for its cytoplasmic retention. Using ten time less AZD8055 (1 μM) for 16 hours, we did not observe any significant cytoplasmic retention of FIE (**Fig. 3D**). Since TOR phosphorylation targets exhibit various sensitivity to TOR activity ^25^, FIE might be a low-sensitive target, only susceptible to (very) strong TOR inhibition, as demonstrated by the notable impact on FIE localization observed after a 3-day treatment with 10 μM of TOR kinase inhibitor ^13^. This robustness suggests a dynamic nature of the TOR-FIE pathway in response to environmental conditions, given that CLF has been established as a regulator of genes in the bistable chromatin state (CS-2 according to Sequeira-Mendes et al., 2009), facilitating rapid stress responses ^14^.

In parallel, we examined whether TOR-CLF pathway influences the transcript levels of H3K27me3-targeted stress-responsive genes. We selected 6 genes within the bistable chromatin context CS-2 (**Fig. 4A**) and 6 genes in the repressive chromatin state CS-5 (**Fig. 4B**) for analysis. Notably, *WRKY70, JAZ1*, and *FMO1* were previously reported to exhibit decreased H3K27me3 deposition upon TOR inhibition ^14^. The transcript levels of these 12 genes were assessed in both *clf-29* CLF-gDNA and *clf-50* CLF-CDS lines, in comparison to their respective wild types (**Fig. 4**). In general, TOR inhibition resulted in the induction of these genes. However, this induction was either abolished or diminished in the *clf*-50 CLF-CDS line for 2 out of the 6 CS-2 genes and for all CS-5 genes. Considering their bistable state, it is noteworthy that the transcriptional regulation of CS-2 genes may require additional mechanisms beyond the TOR-CLF pathway. Moreover, the TOR-FIE dataset revealed diverse expression patterns among our 12 genes ^13^. Some genes exhibited differential regulation in the *fie*-amiR line and the TOR-FIE pathway knockout line (FIE-GFP SSSS/AAAA-line) (**Supplementary figure 3**). Half of these genes were not regulated by FIE or the TOR-FIE pathway. Collectively, our findings reveal a specific role of the TOR-CLF pathway in regulating stress response compared to the TOR-FIE pathway.

**Figure 4:**
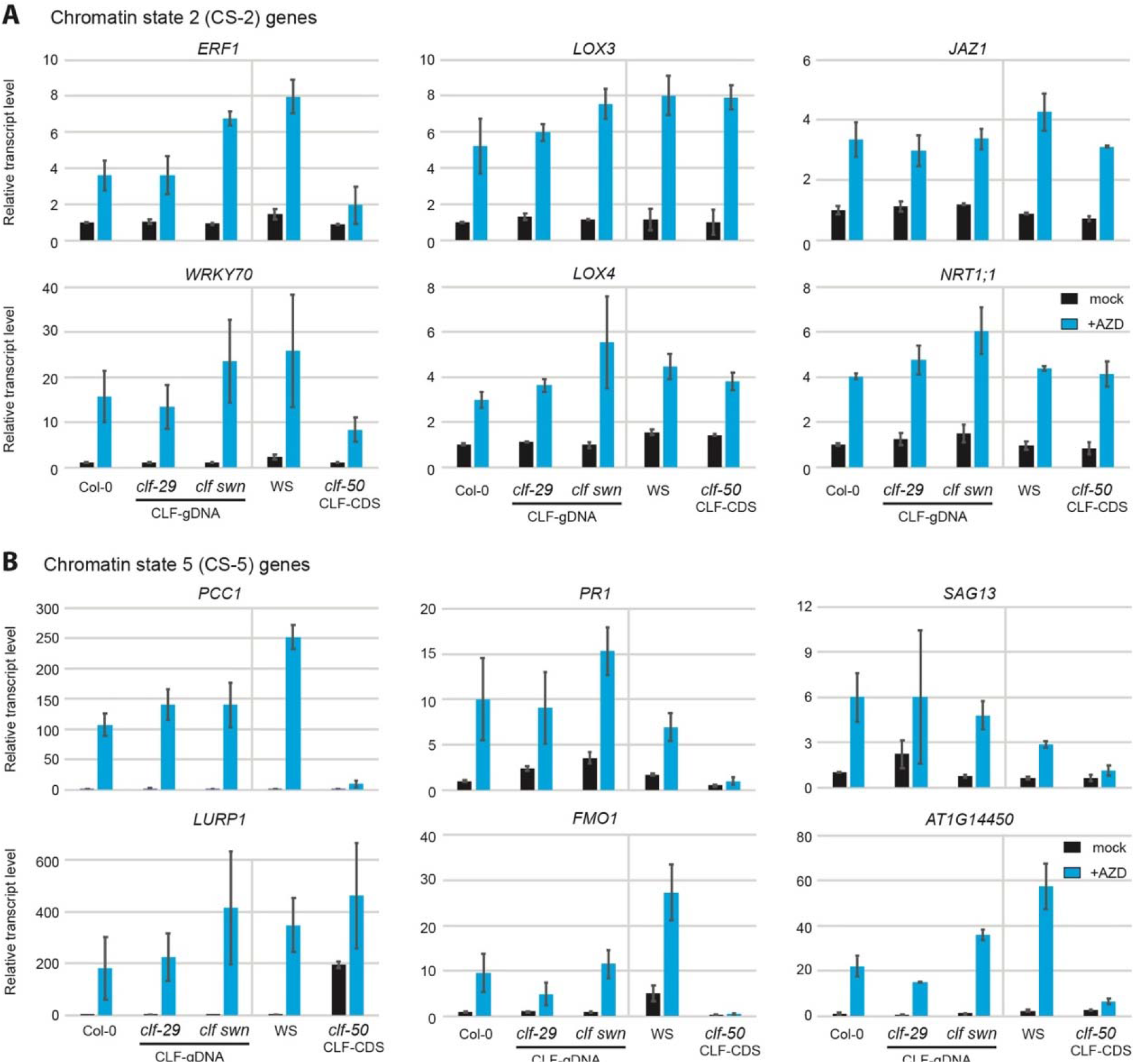
CLF 5’UTR regulates stress-responsive gene expression upon TOR inhibition. Arabidopsis seedlings were treated with 1 μM AZD8055 for 16 h. The relative transcript levels of CS-2 (A) and CS-5 (B) genes was determined in *clf* mutants complemented with either the *CLF* genomic sequence (*clf-29* CLF-gDNA) or coding domain sequence (*clf-50* CLF-CDS) upon TOR inhibition by AZD8055 and expressed relative to Col-0 (n=3, mean±s.d.).

### An evolutionary perspective of TOR-CLF pathway in plant stress response

We sought to determine whether the TOR-uORF regulatory mechanism of H3K27me3-related factors is conserved across species. In stress-sensitive species such as cucumber and cotton, we revealed a consistent pattern of uORFs in *CLF* and *SWN* as in Arabidopsis (**Fig. 5A**). Conversely, rice, known for its submergence tolerance, or the extremophyte *Eutrema salsugineum* exhibited a uORF-less 5’UTR in *CLF* (**Fig. 5B**). Furthermore, the analysis of the *CLF* 5’UTR region in another extremophytes closely related to Arabidopsis, *Crucihimalaya himalaica*, revealed high sequence homology with few SNPs leading to a weak initiation context of AUG from the two uORFs and even shortening of the second uORF (**Fig. 5C**). These findings suggest a dichotomy in the TOR-uORF regulatory mechanism between tolerant and sensitive plant species. The presence or absence of uORFs in *CLF* across different plant species may play a crucial role in shaping their adaptability to stress, influencing stress tolerance mechanisms. Given the potential significance of the TOR-CLF pathway in stress tolerance, particularly in response to *Botrytis cinerea* infection ^14^, we conducted tests using the *clf-50* CLF-CDS and the *clf-29* CLF-gDNA complemented lines. The *clf-50* CLF-CDS line, lacking uORFs in *CLF* and therefore considered as constitutively expressing CLF protein due to impaired TOR regulation, exhibited hypersensitivity to *B. cinerea* infection (**Fig. 5D**). Future investigations into the functional implications of these uORFs in stress response pathways could provide valuable insights into the adaptation strategies employed by various plant species in coping with diverse environmental challenges. These findings underscore the complexity of stress tolerance mechanisms and highlight the need for further exploration to unravel the intricacies of plant stress responses.

**Figure 5:**
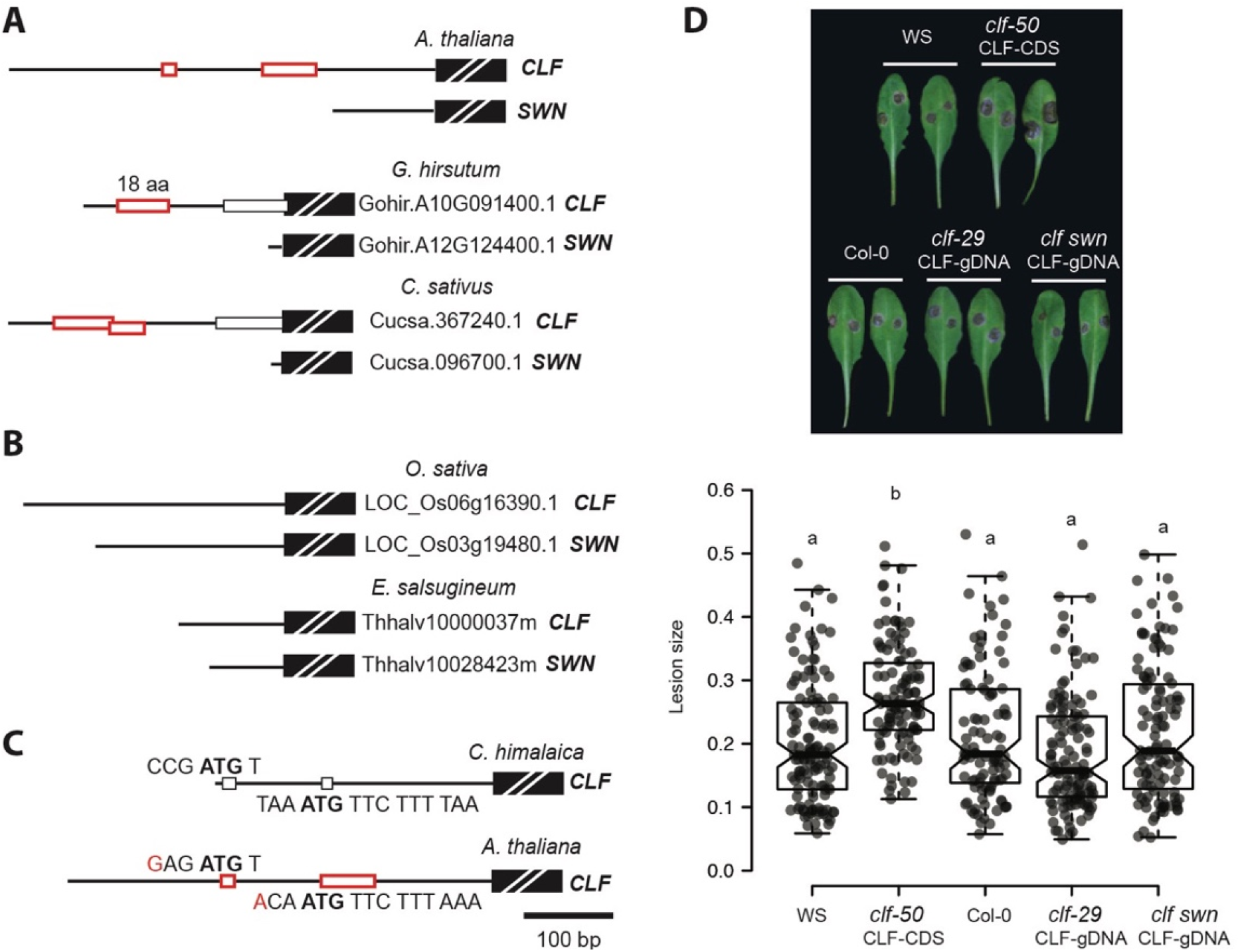
An evolutionary perspective of the TOR-CLF pathway in response to stress conditions. (A-D) Schematic representation of 5 ‘UTR features (black lines) from CLF and SWN across different species, including *A. thaliana* (A), *G. hirsutum* (A), *C. sativus* (A), *E. salsugineum* (B), *O. sativa* (B), and *C. himalaica* (C). Red boxes represent putative inhibitory uORFs with optimal AUG initiation contexts. All 5 ‘UTR sequences were obtained from Phytozome or NCBI. (D) Leaf phenotypes and lesion sizes of WT and *clf* complementation lines infected with *B. cinerea* (n>50, central bars of the notched box represent the median, notches indicate 95% confidence interval, *, *p*<0.05, One-way ANOVA).

## Discussion

In this study, we shed light on the regulatory mechanism behind the TOR-CLF-PRC2 pathway, focusing on the role of uORFs in fine-tuning gene expression. In plants, the 5’UTR of the *CLF* gene harbors a repressive uORF (**Fig. 1A**). Typically, ribosomes translating the uORF dissociate after encountering its stop codon, preventing them from re-initiating translation at the downstream main ORF’s AUG codon. Consequently, the uORF acts as a negative regulator of the main ORF’s translation. In plants, TOR is a well-established master regulator that promotes translation reinitiation following uORF translation through phosphorylation of specific factors such as eIF3h, RISP, and eS6 ^11, 12^. Through this mechanism, TOR can fine-tune CLF protein levels and indirectly influence global H3K27me3 levels in response to environmental cues, independently of SWN, another H3K27 tri-methyltransferase (**Fig. 1**).

The precise mechanisms of re-initiation regulation likely vary depending on the specific functions of the targeted proteins. For instance, TOR-mediated phosphorylation of eIF3h interacts with the ribosome-associated protein RACK1 ^26^, potentially prolonging the association of the 40S ribosomal subunit with initiation factors and aiding in efficient re-initiation ^27^. Interestingly, we showed that eIF3h is required for an efficient translation of CLF (**Fig. 2**). Additionally, phosphorylation of RISP and eS6 by TOR may prevent premature dissociation of the 60S ribosomal subunit from the 40S subunit following uORF translation termination, particularly crucial for re-initiation between closely spaced uORFs within the same mRNA ^11^. These distinct re-initiation control mechanisms mediated by TOR phosphorylation introduce layers of regulatory complexity downstream of TOR signaling pathway. Understanding how these mechanisms integrate with other regulatory pathways could provide deeper insights into plant developmental processes and environmental responses.

Recent studies have expanded our understanding of TOR’s regulatory scope beyond uORFs. For instance, TOR also regulates a specific set of mRNAs containing 5’ oligopyrimidine tract motifs (5’TOPs) through La-related protein 1 (LARP1) phosphorylation, largely impacting genes involved in chromatin regulation such as *LHP1* ^28^. Interestingly, we previously proposed that TOR may regulate H3K27me3 depending on CLF and LHP1 ^14^. To investigate the dynamics of LHP1 translation in response to TOR inhibition, we analyzed *LHP1* mRNA associated with polysomes as above mentioned for *CLF* and *SWN* (**Fig. 1E**). However, we did not observe significant changes in polysome-associated LHP1 mRNA upon short-term TOR inhibition (**Supplementary Fig. 4**), highlighting the context-dependent nature of TOR-LARP1 interactions. Indeed, Scarpin et al., (2020) used seedlings grown under limited photosynthesis in liquid culture and stimulated with glucose. Similar conclusions can be drawn regarding FIE phosphorylation ^13^, especially because we did not observe significant cytoplasmic retention of FIE using less drastic TOR inhibition conditions (**Fig. 3D**). Together, this suggests differential sensitivities and dynamics of TOR targets in response to environmental cues and developmental stages. Moreover besides the conserved FIE phosphorylation site from plants to animals ^13^, CLF uORFs exhibited variations across various plant species adapted to diverse environmental conditions (**Fig. 5A**). This suggests that the response of FIE localization to TOR inhibition may be less dynamic and more specific to developmental transitions, while the uORF-mediated CLF translational control represents a finetuned response towards fluctuating environmental conditions. Because such evolutionary adaptations in uORF sequences indicate diversification in response to environmental stimuli, investigating how they affect TOR-mediated regulation of translation initiation appears crucial for understanding plant adaptation strategies.

Despite the identification of numerous TOR targets in plants, the sensitivity and dynamic responses of these targets to TOR activity remain a significant challenge. Our study, together with previous findings ^13, 14, 28^, supports that TOR phosphorylates three distinct protein sets to regulate H3K27me3: LARP1 for control of LHP1 translation, eIF3h for CLF translation, and FIE for PRC2 assembly. These findings open up exciting avenues for future research aiming to decipher the specific developmental contexts in which TOR signaling pathways operate and how they coordinate with each other. This integrated approach will provide a more comprehensive understanding of TOR signaling in plants and its implications for growth, development, and stress responses.

## Methods

### Plant materials and growth conditions

Different complementation lines, including *fie* FIE-GFP, *clf-29* CLF-gDNA, *clf swn* CLF-gDNA and *clf-50* CLF-CDS lines, were kindly provided by Dr. Francois Roudier (CNRS, France) ^23, 24^. *Arabidopsis thaliana* Col-0 was used as the wild-type model in this study, while the ecotype WS served as the wild-type control for the *clf-50* CLF-CDS line. All seedlings used in this study were germinated and grown on ½ MS medium supplemented with 0.5% sucrose and 1.2% agar in a growth chamber with a 16-hour day/8-hour night cycle. To analyze growth under TOR inhibition conditions, 0.2 μM or 0.5 μM AZD8055 was added to the ½ MS medium. Short-term TOR inhibition was conducted in liquid ½ MS medium with 1 μM AZD8055 for 16 hours. CaMV infection was performed according to a previously described method for 22 days ^11^.

### GUS assay in Arabidopsis protoplasts

GUS assay was performed as described previously ^29^. Arabidopsis protoplasts prepared from 1-week-old seedlings (mesophyll protoplasts) were transfected with plasmid DNA by the PEG method. Standard short 5’UTR pmonoGUS was used as basal translation control. The constructs of native or mutated CLF 5’UTR fused to GUS (5’*CLF*-GUS and 5’*CLF(ATG to TAG)*-GUS) were generated by substitution of a short leader sequence in pmonoGUS plasmid with the native or mutated CLF 5’UTR, downstream of the constitutive 35S promoter and upstream of the start codon of the GUS reporter using HiFi DNA assembly cloning kit (NEB E5520). Fragments of 35S promoter + vector backbone + GUS was amplified using forward primer ATGTTACGTCCTGTAGAAACCC and reverse primer GTCGACTCCAAATGAAATGAAC. 5’UTR of CLF with flanking sequence used for assembly was amplified with forward primer GTTCATTTCATTTGGAGTCGACAAAAAAAAAACAAAAAATAAAGCCCAAAC and reverse primer GGGTTTCTACAGGACGTAACATTGTCAAGAAACCAGATCGGAAC. 2^nd^ uORF was mutated using two additional complementary primers CCAATTCGTAAACATTGTTCTTTAAAAG and CTTTTAAAGAACAATGTTTACGAATTGG. After overnight incubation (16-18 h), transfected protoplasts were harvested by centrifugation and resuspended in GUS extraction buffer (50 mM NaH2PO4 pH 7.0, 10 mM EDTA, 0.1% NP-40). The aliquots were immediately taken for GUS assays and RNA extraction. GUS activity was measured in 5, 10 and 20 min by FLUOstar OPTIMA fluorimeter (BMG Biotech).

### Polysome profiling

200 mg 10 days old seedlings were homogenized in 600 μL extraction buffer (200 mM Tris-HCl pH9.0, 200 mM KCl, 25 mM EGTA pH 8.0, 35 mM MgCl2, 1 % DOC, 1 % PTE, 1 % BriJ-35, 1 % Triton-X100, 1 % Igepal, 1 % Tween-20, 10 mM DTT, 10 μM Mg132, protease inhibitor EDTA-free) by rotation at 4°C for 20 minutes. After centrifugation at 16,000 g for 10 minutes at 4°C, 400 μL of the extract were loaded on the top of a 7-47 % sucrose gradient and centrifuged in a SW41 rotor (Beckman) for 3h at 38 000 rpm at 4°C. After centrifugation, the gradients were analyzed with absorbance at 254 nm and fractioned. Different fractions were mixed with 1 volume of 8 M Guanidine-HCl and 2 volume of 70 % ethanol and stored over night at -20°C. RNA was precipitated by centrifuge at max speed at 4°C for 45 min. The quality of precipitated RNA was monitored by 1% agarose gel and used further for cDNA synthesis and qPCR. RNA was extracted with Trizol and converted to cDNA using the RevertAid H Minus First Strand cDNA Synthesis Kit (Thermo Scientific). Primers are listed in Supplementary Table S1.

### Western blot

For immunological detection, total soluble proteins were extracted from 50 mg plant materials with 250 μL 2x Laemmli buffer. Proteins were denatured for 5 min at 95°C and separated on 15% SDS-PAGE. Subsequently, proteins were blotted to PVDF membrane. The primary antibodies anti-GFP (1:3000, Invitrogen, A-11122), anti-H3K27me3 (1:5000, Agrisera, AS16-3193) and anti-H3 (1:5000, Agrisera, AS10-710) were detected using the HRP-conjugated secondary antibody (1:20,000).

### RT-qPCR

RNA was extracted with Trizol and converted to cDNA using the RevertAid H Minus First Strand cDNA Synthesis Kit (Thermoscientific). Transcript abundance was determined using gene-specific primers listed in Supplementary Table S1 on a LightCycler 480 Real-time PCR system (Roche Diagnostics) in a final volume of 10 μL SYBR Green Master mix (Eurogentec). AT4G26410 (RGS1-HXK1 INTERACTING PROTEIN 1, RHIP1), AT1G13440 (GLYCERALDEHYDE-3-PHOSPHATE DEHYDROGENASE, GAPC2) and AT4G34270 (INTERACTING PROTEIN OF 41 KDA, TIP41) were used as internal references. Relative expression values were calculated using the comparative cycle threshold method 2^-ΔΔCt^.

### Fungi infection

?Five-week-old, soil-grown plants were inoculated with *Botrytis cinerea* in a short-day chamber (12 hours light/12 hours dark, with light intensity of 80-100 μmol m ^−2^s ^−1^, 22°C day/18°C night temperature, and 50% relative humidity). For the lesion assay, 5 μL droplets of a *B. cinerea* spore suspension were directly applied to the upper leaf surface. Following inoculation, plants were maintained at high humidity to promote infection. Lesion size was measured three days after inoculation.

### Microscope analysis and quantification

Seedlings were germinated and grown for 7 days on ½ MS agar vertical plates. They were then transferred to ½ MS liquid medium supplemented with 0.5% sucrose for 24h before being treated with 1 μM AZD8055 for 16 hours. Following treatment, seedlings were mounted on microscope slides in water and examined using a Zeiss LSM 700 microscope equipped with 20x/0.8 NA lens. The excitation and emission wavelengths for the fluorescent protein GFP were 488 nm and 510 nm, respectively. Confocal images were acquired from the epidermal and cortex layers of the root of at least 6 distinct seedlings per genotype and fluorescent signals were measured on nuclei (n≥196) using ImageJ.

## Supporting information

Supplementary informations

## Acknowledgment

We thank Dr. Francois Roudier (ENS-RDP, France) for sharing the complementation lines of *fie, clf* and *clf swn* mutants, Dr. Thierry Heitz (IBMP-CNRS, France) for sharing the *B. cinerea* culture, Dr. Csaba Papdi, Dr. Lyuba Ryabova (IBMP-CNRS, France), Dr. Christel Carles (LPCV, France) for stimulating discussions during the course of this research project.

## Author contribution

Y. D. and A. B. initiated and coordinated the project. Y. D., V. V. U. and A. B. designed the experiments. Y. D., E-Z. O., E. H., W. Z., and A. B. performed experiments. Y. D. and A. B. made figures and wrote the manuscript. V. V. U. revised the manuscript. All authors approved the final version of the manuscript.

