## Supplementary material for "Target of rapamycin (TOR) regulates *CURLY LEAF* (*CLF*) translation in response to environmental stimuli": Table S1

| Name | forward | reverse | Gene ID | Details |
|---|---|---|---|---|
| CLF 5'UTR | CGAGCTACGTGATATTCCGGT | GAAAATAGCGACGTGGCAGC | AT2G23380 | |
| SWN 5'UTR | TTCCAGTCCACACTCCGTTC | GTTGCTATCGTCCGTCACCA | AT4G02020 | |
| CLF CDS | TCGCTAAAGAAGAAGCTTGCAG | ACAGCTACCTCCTCGTTCCA | AT2G23380 | |
| SWN CDS | CGCTAAGGTGATGTTTGTAGCAG | CTTGGAGCCTTCAGGTTTGC | AT4G02020 | |
| PCC1 | ACAAACTCCAAGGGCGTCAA | GATGTACAGAGGCTGGAGCA | AT3G22231 | state 5 gene upregulated in clf28 |
| LURP1 | GTACGGAGGAGGGTGCTCTA | TTCCGAAGAAAAAGCCTCGC | AT2G14560 | state 5 gene upregulated in both clf28 and clf29 |
| PR1 | CTCGGAGCTACGCAGAACAA | CGCTACCCCAGGCTAAGTTT | AT2G14610 | state 5 gene upregulated in both clf28 and clf29 |
| FMO1 | GCTTGAGTTTCCAAGCGGTG | CGGCTTAGCCACATTGAACG | AT1G19250 | state 5 gene upregulated in clf28 |
| SAG13 | CATCGGGGAAGCTGTGGT | TCGCAGACAGAAGTGGTGAC | AT2G299350 | state 5 gene upregulated in both clf28 and clf29 |
| AT1G14450 | GGGCAGTGAAAGTCGGAAGA | TCCCGGGAGTTCACCACTAT | AT1G14450 | state 5 gene targeted by CLF |
| ERF1 | TCTAATCGAGCAGTCCACGC | CTCTTATCTCCGCCGCGAAT | AT3G23240 | state 2 gene targeted BRM |
| WRKY70 | TGGTTCGTCCACGGAGAATG | CCCATTGACGTAACTGGCCT | AT3G56400 | state 2 gene targeted BRM |
| JAZ1 | CTCGTGAAGGAGGGCAAACT | CGGGAATGTGTCTCGGTTCA | AT1G19180 | state 2 gene targeted BRM |
| LOX3 | CACACTTAAGCCGGTAGCCA | ACTGGAGGTGTAAGCACACG | AT1G17420 | state 2 gene targeted BRM |
| LOX4 | CCTAGCCGTAGGAATCGCTG | CACGTAACACCCGGTTCAGA | AT1G72520 | state 2 gene targeted BRM and upregulated in clf28 and clf29 |
| NRT1;1 | TAAGGGATCAGGAAGCGGGA | AAGAGGATGCATGTTGCCCA | AT1G12110 | state 2 gene targeted BRM |
| GAPDH | TTGGTGACAACAGGTCAAGCA | AAACTTGTCGCTCAATGCAATC | AT1G13440 | |
| EXP | GAGCTGAAGTGGCTTCAATGAC | GGTCCGACATACCCATGATCC | AT4G26410 | |
| TIP41 | GTGAAAACTGTTGGAGAGAAGCAA | TCAACTGGATACCCTTTCGCA | AT4G34270 | |
