## Supplementary informations for "Target of rapamycin (TOR) regulates *CURLY LEAF* (*CLF*) translation in response to environmental stimuli"

**5’UTR sequence of CLF**

AAAAAAAAAACAAAAAATAAAGCCCAAACAAATAGGGTTTAAGAGCCCATTAA
ATCAGAGCCCAGCAACGTTCGTGTTTGTTTTGTTCAGTTGATATAGAACCAGTCTC
GAACTCTAGTAGATTCGTTGTCTCGAGCTACGTGATATTCCGGTAAACCGTCGCC
GGAGATGTACAGTAGATAATTCAAATCGAGCTGCCACGTCGCTATTTTCCCACCA
ACTTTTTTTTTTTTAAATAATTGGAAAAAGTTTCTAATTTACACGCTTCCCAATTCG
TAAACAATGTTCTTTAAAAGTAAAACGAAATTTTTTTTTGGATTTGAAGATAAAAG
ATCCAAAATAAAAACAAAATTGAAGATAAATTTTTAGGGTTTATAACGACCCGCC
AAATTCCTTCTCTCCTCCAACATTTCAGATTTCGCGACCCGGATCTTAACCCGGAC
CCGCATTTGTTTCGGTTCCGATCTGGTTTCTTGACAATG...

**5’UTR sequence of CLF in WS**

AAAAAAAAAACAAAAAATAAAGCCCAAACAAATAGGGTTTAAGAGCCCATTAA
ATCAGAGCCCAGCAACGTTCGTATTTGTTTTGTTCAGTTGAAATAGAACCAGTCT
CGAAATCTAGTAGATTCATTGTCTCGAGCTACGTGATATTCCGGTAAACCGTCGC
CGGAGATGTACAGTAGATAATTCAAATCGAGCTGCCACGTCGCTATTTTCCCACC
AACTTTTTTTTTATTAAATAATTGGAAAAAGTTTCTAATTTACACGCTTCCCAATT
CGTAAACAATGTTCTTTAAAAGTAAAACGAAATTTTTTTTTGGATTTGAAGATAAA
AGATCCAAAATAAAAACAAAATTGAAGATAAATTTTTAGGGTTTATAACGACCCG
CCAAATTCCTTCTCTCCTCCAACATTTCAGATTTCGCGACCCGGATCTTAACCCGG
ACCCGCATTTGTTTCGGTTCCGATCTGGTTTCTTGACAATG...

**Figure S1. 5’UTR sequences of *CLF* genes in Col-0 and WS ecotypes.** ATG codon are in red, with the codon for uORFs underlined. Two uORFs are highlighted in yellow with in-frame stop codon in blue. SNPs in WS are highlighted in green.

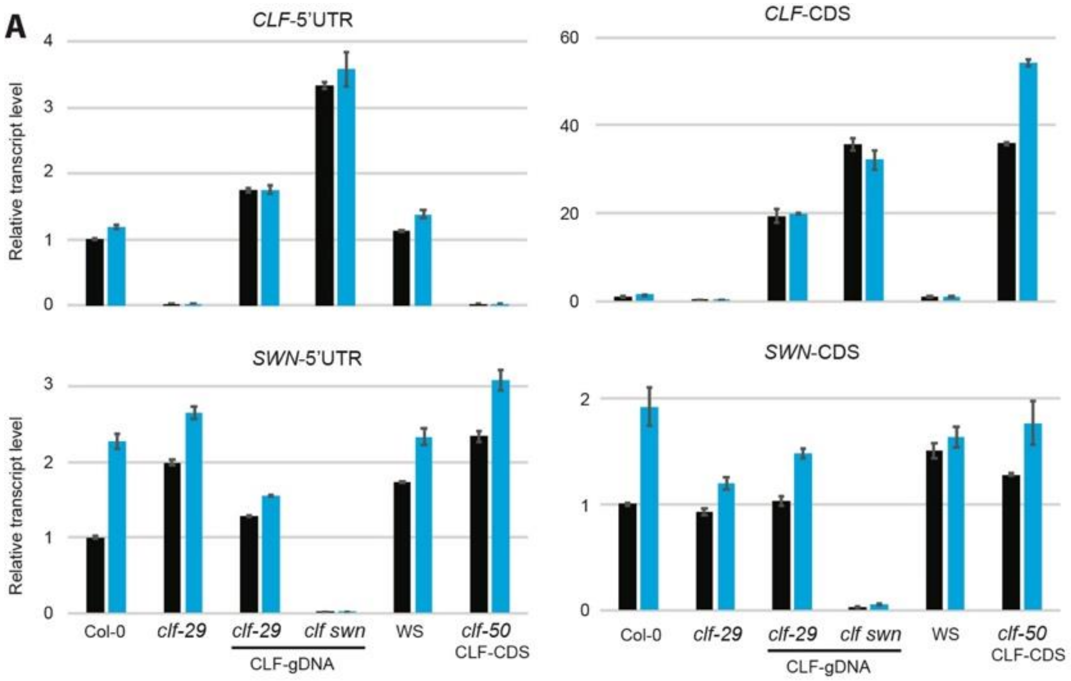


**Figure S2. Relative transcript levels of *CLF*, and *SWN* upon TOR inhibition.** Arabidopsis seedlings were treated with 1 µM AZD8055 for 16 h. The relative transcript level of *CLF*, and *SWN* was determined in clf mutants complemented with CLF genomic sequence (*clf-29* CLF-gDNA) or coding domain sequence (*clf-50* CLF-CDS). The *clf-29* and *swn-4* (in the background of *clf-29* CLF-gDNA) mutants were used as controls. For *CLF* and *SWN*, specific primer pairs were designed to target both 5'UTR and CDS (n = 3, mean ± s.d.).

| | *fie*-amiR | FIE AAAA | TOR | TOR-CLF |
|---|---|---|---|---|
| ***PCC1*** | -1.354 | 1.565 | Y | Y |
| ***LURP1*** | 0.000 | 6.419 | Y | Y |
| ***PR1*** | 0.000 | 4.768 | Y | Y |
| ***FMO1*** | 0.000 | 0.000 | Y | Y |
| ***SAG13*** | 0.000 | 4.450 | Y | Y |
| ***AT1G14450*** | 0.000 | 0.000 | Y | Y |
| ***ERF1*** | 0.000 | 0.000 | Y | Y |
| ***WRKY70*** | -2.267 | -0.098 | Y | Y |
| ***JAZ1*** | 0.000 | 0.000 | Y | N |
| ***LOX3*** | 0.000 | 0.000 | Y | N |
| ***LOX4*** | 1.838 | -1.507 | Y | N |
| ***NRT1;1*** | 0.000 | 0.000 | Y | N |

**Figure S3: Stress-responsive gene expression affected by TOR, FIE and CLF.** CS-2/5 gene expression was analysed from several datasets. Numbers indicate log2(fold change) in FIE amiRNA silencing line (fie-amiR) and the TOR-FIE pathway knockout line (FIE AAAA) obtained by Ye et al. (2022). "Y" denotes their expression is regulated by TOR (induced by AZD as shown in Fig. 4) and TOR-CLF pathway (impaired induction by AZD in *clf50* CLF-CDS line as shown in Fig. 4). "N" indicates that their expression is not regulated by the TOR-CLF pathway.

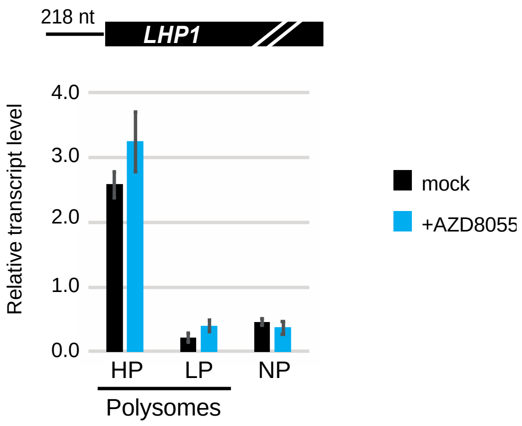


**Figure S4: LHP1 translation efficiency upon TOR inhibition.** Polysome profiling was performed using 7-day-old seedlings treated with 1 µM AZD8055 for 16 hours. *LHP1* mRNA levels was quantified in heavy polysome (HP), light polysome (LP) and non-polysome (NP, including 80S, 60S and 40S fractions) fractions (n = 3, mean ± s.d., $^{*}p < 0.05$).
